# Video Foundation Models for Animal Behavior Analysis

**DOI:** 10.1101/2024.07.30.605655

**Authors:** Jennifer J. Sun, Hao Zhou, Long Zhao, Liangzhe Yuan, Bryan Seybold, David Hendon, Florian Schroff, David A. Ross, Hartwig Adam, Bo Hu, Ting Liu

**Affiliations:** Google. Currently at Google DeepMind

**Author notes:** Equal senior contribution (Bo Hu, Ting Liu).

**Keywords:** video analysis, animal behavior, foundation models, behavior classification

## Abstract

Computational approaches leveraging computer vision and machine learning have transformed the quantification of animal behavior from video. However, existing methods often rely on task-specific features or models, which struggle to generalize across diverse datasets and tasks. Recent advances in machine learning, particularly the emergence of vision foundation models, i.e., large-scale models pre-trained on massive, diverse visual repositories, offers a way to tackle these challenges. Here, we investigate the potential of *frozen* video foundation models across a range of behavior analysis tasks, including classification, retrieval, and localization. We use a single, frozen model to extract general-purpose representations from video data, and perform extensive evaluations on diverse open-sourced animal behavior datasets. Our results demonstrate that features with minimal adaptation from foundation models achieve competitive performance compared to existing methods specifically designed for each dataset, across species, behaviors, and experimental contexts. This highlights the potential of frozen video foundation models as a powerful and accessible backbone for automated behavior analysis, with the ability to accelerate research across diverse fields from neuroscience, to ethology, and to ecology.

## 1 Introduction

The proliferation of video recording technology has revolutionized the study of naturalistic animal behavior, enabling researchers to capture vast amounts of behavioral videos in unprecedented detail [1]. However, extracting meaningful insights from these large-scale video datasets is challenging – manual annotation by experts to accurately quantify complex behaviors is both time-consuming and expensive, and does not scale well to the rapidly growing volume of video data. Additionally, the specific research questions and target behaviors vary significantly across studies, further complicating the development of a unified video analysis model for animal behavior.

Foundation models [2] represent a recent paradigm shift in machine learning, and have already demonstrated remarkable generalization capabilities across diverse domains [3, 4]. These large-scale models are pre-trained on massive, heterogeneous datasets, enabling them to learn a wide range of representations that can be transferred to new tasks with minimal or no model adaptations. This inherent advantage addresses a common bottleneck in computational approaches of animal behavior analysis – generalizability to new domains or tasks (Figure 1).

**Fig. 1.**
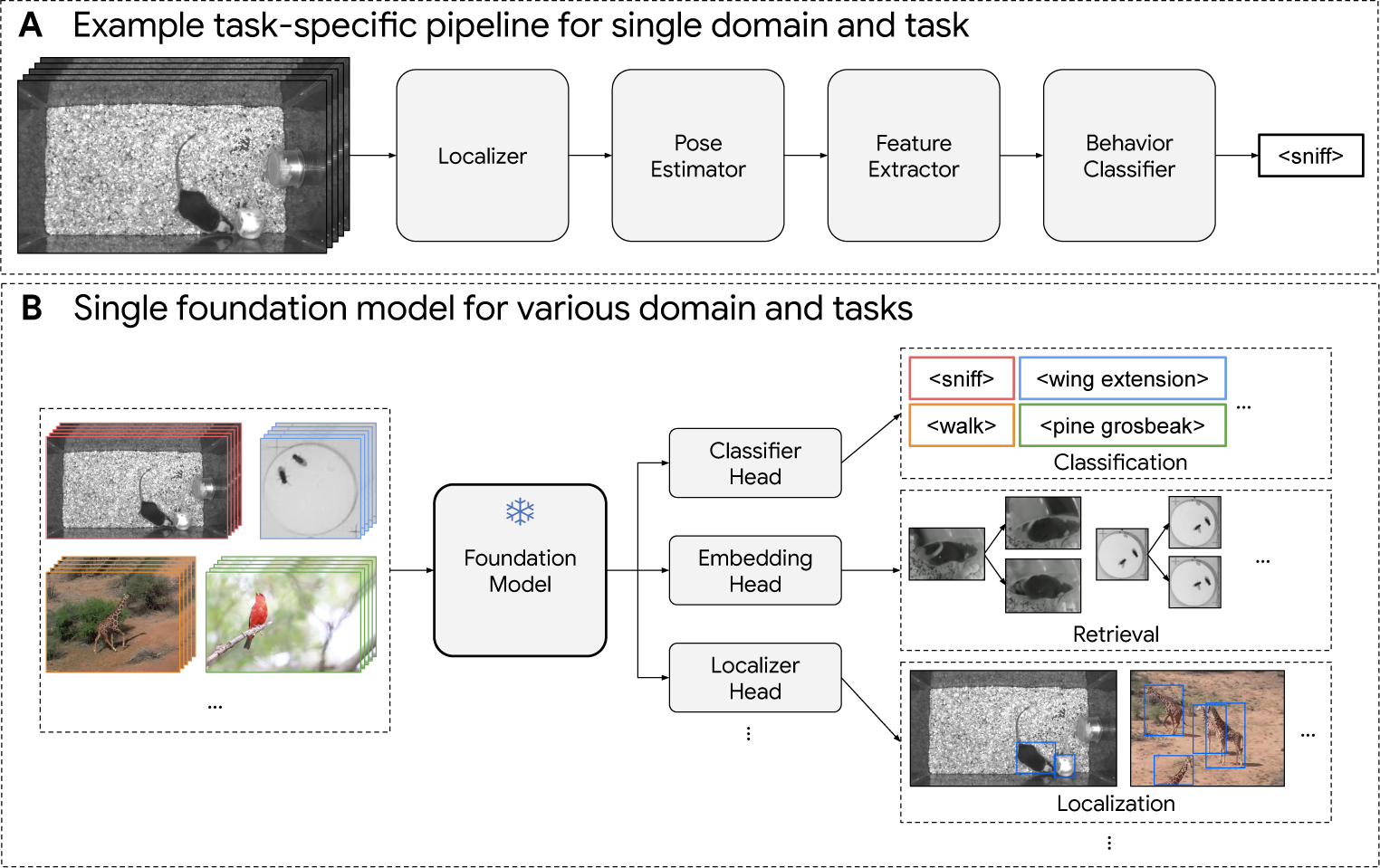
Overview of foundation model for animal behavior analysis. **(A)** Previous work typically require pipelines with specially-designed modules and/or hand-crafted features specific for individual domains and tasks. **(B)** In this work, we leverage recent development in foundation models and demonstrate that a single set of frozen features from the pre-trained general-purpose models, paired with minimal task-specific decoder heads, is able to generalize to a range of various downstream tasks across various animal behavior domains.

While existing computational methods have shown promising results on specific tasks in animal behavior analysis [5–10], these approaches typically rely on models or features that are highly tailored to the task. These features could be hand-crafted (e.g., distance between agents, speed of an agent), learned from the data (e.g. using machine learning), or a combination [5, 7, 8, 11]. Some approaches in this space use pre-trained models (e.g., ResNet [12] trained on ImageNet [13]) for transfer learning [5, 14–16], which still require significant task-specific fine-tuning to adapt the backbone to datasets containing different species or behaviors. In contrast, the core innovation of foundation models lies in their ability to be used as a single, frozen backbone, with fixed weights across diverse tasks and experiments. The use of a unified set of frozen features for behavior analysis has the potential to streamline task-specific pipelines, and reduce the need for individual fine-tuning.

Here, we present the first investigation of frozen video foundation models on a wide range of animal behavior analysis tasks, including classification, retrieval, and localization. In a significant departure from conventional approaches that rely on extensive model adaptation, we utilize a single, frozen video foundation model (VideoPrism [17]) to extract general-purpose representations. This model was pre-trained on large-scale general-domain Internet data, with no additional training on animal-specific videos. We aim to study the effectiveness of using general visual representations learned from Internet data to tasks important in animal behavior analysis.

Our evaluation on diverse open-sourced animal behavior datasets reveals the capability of video foundation models to generalize across species, behaviors, and experimental paradigms, with minimal or no decoder training. We focus on datasets captured as part of real-world scientific experiments in fields such as neuroscience, ethology, and ecology. While the majority of datasets in our evaluation feature mice [5, 9, 18], a common model organism in behavioral research with many largescale open-sourced video datasets, we also assess performance on other species such as flies [19], birds [20], giraffes, and zebras [21] when there are available, annotated video datasets. We compare the video foundation model to existing baselines, including domain-specific models designed by experts and trained specifically for each dataset, as well as a frozen image foundation model (CLIP [4]).

Results demonstrate that frozen features from video foundation models achieve better performance compared to image models, and competitive performance compared to task-specific baselines, when expert-designed baselines are available. This suggests that a unified video foundation backbone could potentially reduce the need for time-consuming and costly development of task-specific pipelines. Ultimately, our finding paves the way for a more interconnected computational framework that leverages state-of-the-art machine learning and computer vision tools for studying animal behavior, accelerating scientific discovery and promoting collaboration across disciplines.

### Background on foundation models

Thanks to the recent success in large language models [22, 23], wide research interests in the fields of machine learning and computer vision are shifting from designing domain-specific models to building large foundation models. Most recent foundation models in the vision domain focus on images [24–28]. These models employ transformers specifically trained to align web-scale image-text paired data through contrastive learning [4, 29], predicting the next token in the language modality [30] or in the interleaved image-text sequence [31]. Recent models are also trained on a mixture of these objectives [32, 33]. While some of these foundation models are able to take video frames as input, they fall short on motion and temporal modeling [34]. In this paper, we consider CLIP [4] as the representative image foundation model for evaluation due to its strong performances on a wide range of vision tasks [35–37].

Compared with image inputs, videos contain additional temporal dynamics, and hence are more challenging to model. Existing video foundation models are mainly trained to model temporal information using self-supervised learning over the video-only modality [38–46] or video-language modeling of videos with captions [47–53] (e.g., alt-text or transcribing text from audio). However, a recent study [54] points out that existing video-language models lack knowledge of actions, while self-supervised models from video-only data usually struggle with semantics. In contrast, a recent video foundation model, VideoPrism [17], tackles both of these challenges. It obtains state-of-the-art results over 31 video understanding tasks by producing video representations from one single frozen model. Therefore, we use VideoPrism as the video foundation model in our experiments. We aim to unlock the potential of video foundation models for automated animal behavior analysis and pave the way for the development of more generalizable and effective video understanding models in the future.

## 2 Results

Across ten datasets, we find that VideoPrism outperforms baselines on tasks including video behavior classification, few-shot classification, retrieval, and localization (Figure 2).

**Fig. 2.**
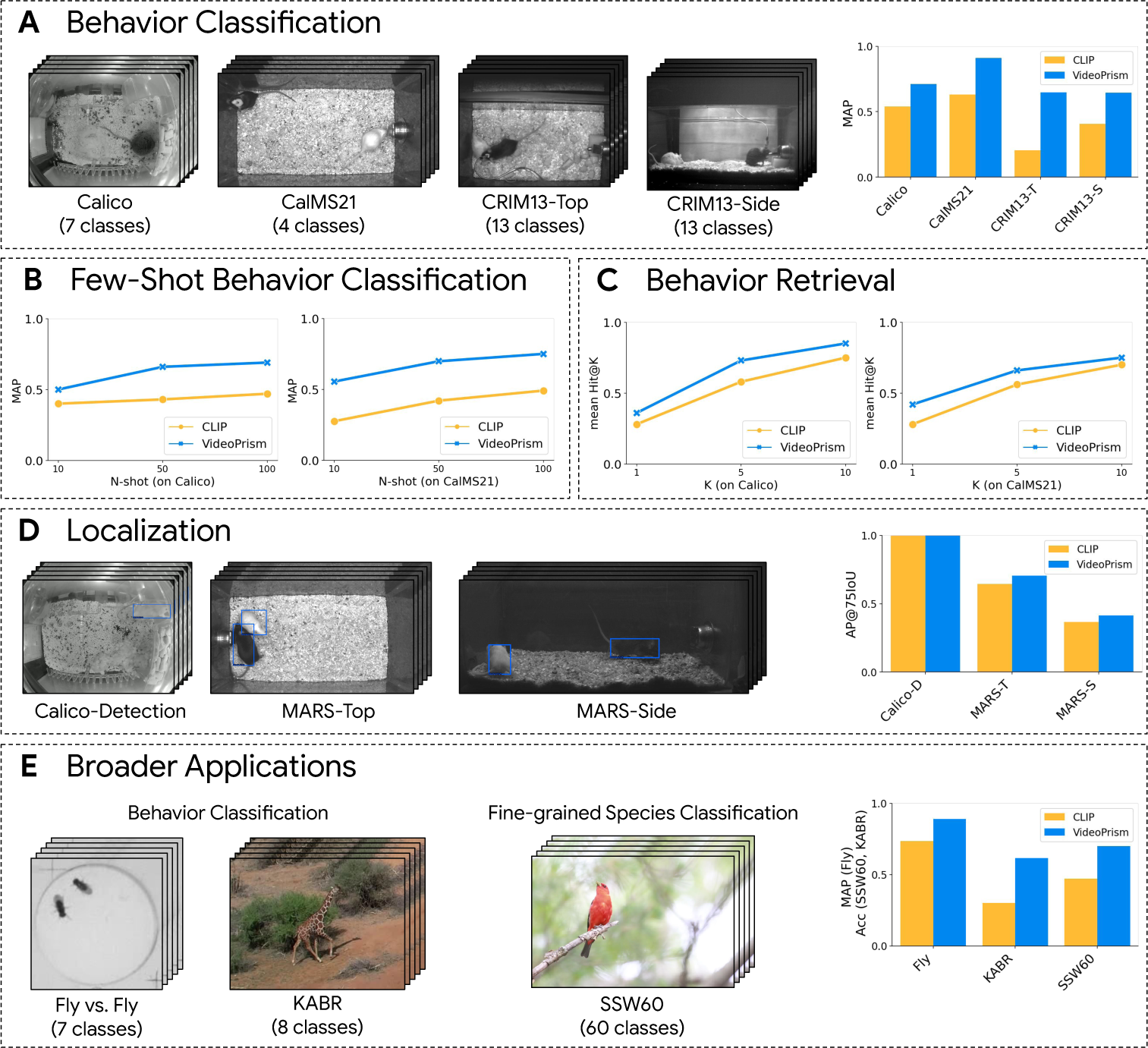
Summary of experiments. We study the performance of image foundation model (CLIP) and video foundation model (VideoPrism) on a wide range of open-sourced animal video benchmarks, including **(A)** behavior classification on four mouse datasets, **(B)** few-shot behavior classification and **(C)** behavior video retrieval on two mouse datasets, **(D)** mouse localization on three mouse datasets, and **(E)** broader applications on fly behavior classification (Fly vs. Fly), Kenyan animal behavior classification from drone videos (KABR), bird species classification (SSW60). Results demonstrate that video foundation models significantly improve performance relative to image models.

We briefly summarize each dataset in Table 1, and compare performance of each method on these datasets. We include VideoPrism (video foundation model), CLIP (image foundation model), as well as specialized models developed by the originally published dataset papers, when available. Notably, the backbone weights of the foundation models are not trained on these datasets, and features from one frozen foundation model is used for all experiments, with either no or minimal decoder adaptations. Our tasks include behavior classification (Section 2.1), few-shot behavior classification (Section 2.2), video-based behavior retrieval (Section 2.3), localization (Section 2.4), as well as broader applications to other organisms and tasks (Section 2.5).

**Table 1.** Summary of datasets. We use open-sourced annotated video datasets released by researchers from fields including neuroscience, ethology, and ecology (see more details in Section 4.5). The number of annotations in the last column either represents one annotation per frame, per clip (segmented from videos), or per video, depending on the expert annotations provided in the original dataset. We use the dataset split published in the original papers, whenever this split is available.

| Results Section | Dataset | Metric | # Annotations (Train/Test) |
| --- | --- | --- | --- |
| Behavior classification | Calico [55] | mAP | 1,253 / 400 clips |
|  | CalMS21 [18] | mAP | 507,660 / 262,107 frames |
|  | CRIM13-Top [9] | mAP | 1,630,114 / 1,994,885 frames |
|  | CRIM13-Side [9] | mAP | 1,612,327 / 1,892,817 frames |
| Few-shot behavior classification | Calico [55] | mAP | (10/50/100)-shot / 400 clips |
|  | CalMS21 [18] | mAP | (10/50/100)-shot / 262,107 frames |
| Behavior retrieval | Calico [55] | mean Hit@K | 1,253 / 400 clips |
|  | CalMS21 [18] | mean Hit@K | 507,660 / 262,107 frames |
| Localization | Calico-Detection [55] | AP@IoU | 5,460 / 2,124 frames |
|  | MARS-Top [5] | AP@IoU | 10,000 / 5,000 frames |
|  | MARS-Side [5] | AP@IoU | 10,000 / 5,000 frames |
| Broader Applications | Fly vs. Fly [19] | mAP | 1,067,329 / 322,393 frames |
|  | KABR [21] | Macro-Acc. | 1,545,513 / 290,021 frames |
|  | SSW60 [20] | Accuracy | 3,462 / 1,938 videos |

### 2.1 Behavior classification

We assess the ability of frozen foundation models to generalize across diverse mouse behavior datasets and tasks (Table 2). Our evaluation spans three mouse datasets: Calico [55], CalMS21 [18], and CRIM13 [9]. Calico and CalMS21 are top-view only, while CRIM13 consists of both top and side view. These datasets encompass a wide range of behaviors, from social interactions in resident-intruder assays to actions in home-cage environments. For each dataset, we compare the performance of domain-specific baselines, developed by experts and trained on each dataset, with frozen image foundation model (CLIP) and video foundation model (VideoPrism). The specialized model for Calico is ResNet with a 2-layer MLP classification head, following [55]. The specialized model for CalMS21 first extracts trajectory features, then uses a 1D-CNN for temporal aggregation [18]. For CRIM13, we conduct evaluations for top-view and side-view separately^1^. For the foundation models, a trainable pooling layer and a linear classifier is used to map the features to the class outputs.

**Table 2.** Behavior classification results. We use mean average precision across all classes. For CalMS21 and CRIM13, we only average over behaviors-of-interest and do not include the background class in metric computation, following [18].

| Method | Calico | CalMS21 | CRIM13 Side | CRIM13 Top |
| --- | --- | --- | --- | --- |
| Specialized models | 52.6 | 88.9 | - | - |
| CLIP | 51.8 | 62.7 | 34.0 | 14.1 |
| VideoPrism | 71.2 | 91.1 | 64.5 | 64.9 |

The video foundation model consistently matches or exceeds the performance of domain-specific models, highlighting its potential to reduce the need for laborious and costly development of task-specific approaches. Additionally, the video foundation model significantly outperforms the image-based model, showing the importance of incorporating temporal information during feature extraction for studying behavior. Visual comparison of VideoPrism predictions and expert annotations on CalMS21 and CRIM13 datasets (Figure 3), both containing frame-level annotated videos, demonstrates the model’s capacity to capture both extended behaviors (e.g., “mount”) and those characterized by rapid transitions (e.g., “attack”). The VideoPrism model misses bouts of certain behaviors (e.g., “walk away”, “approach”) on CRIM13, indicating the room for future improvement, for example, by integrating a larger window of temporal context.

**Fig. 3.**
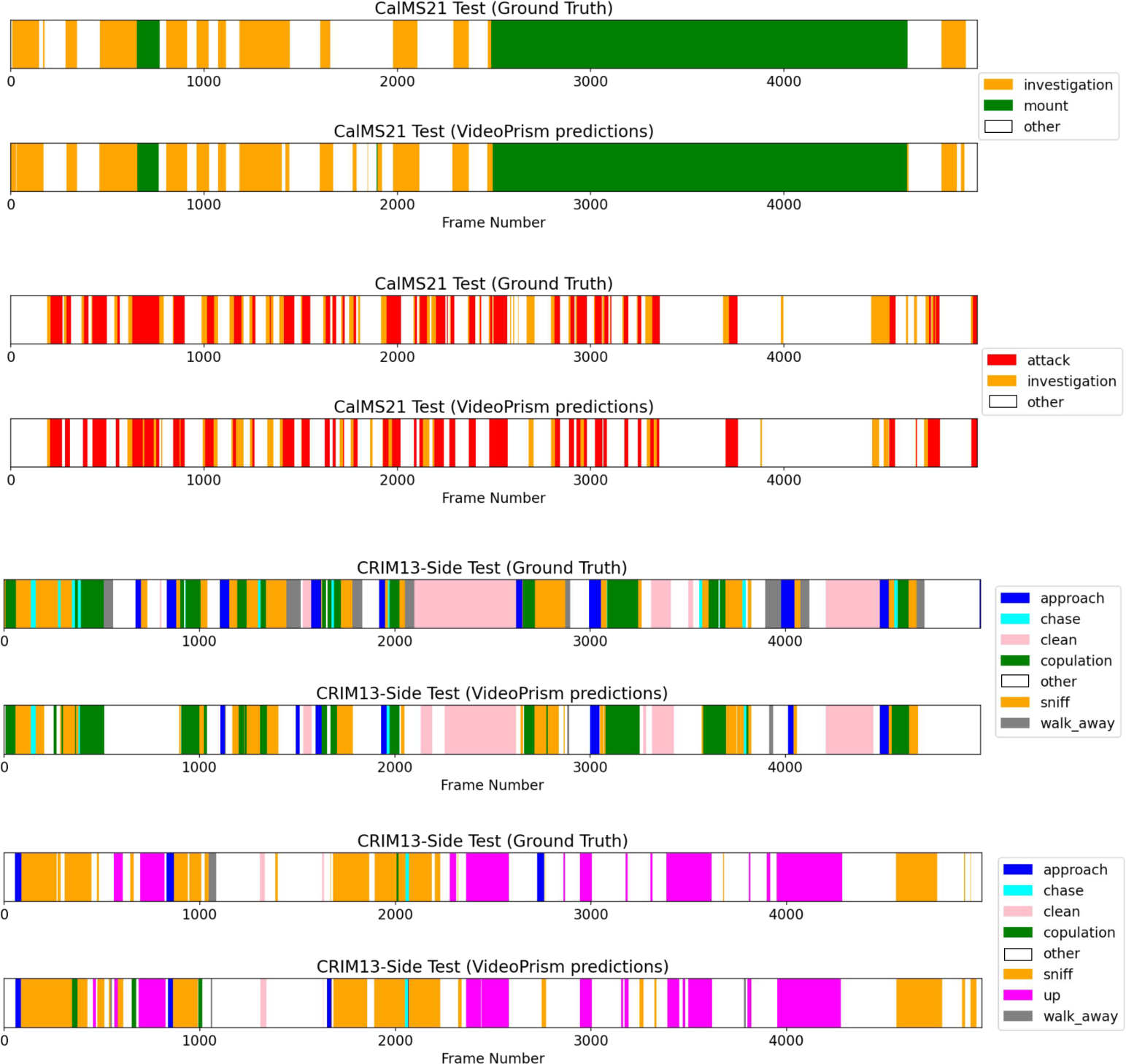
Behavior classification using VideoPrism. We show examples of VideoPrism’s behavior classification performance on CalMS21 and CRIM13. The top strips in each setting show the expert annotations, and the bottom are model predictions. These predictions are obtained by applying the classifier in a 16-frame sliding window, averaging the predictions across all sliding windows applied to a single frame, and taking the class with the maximum likelihood.

### 2.2 Few-shot behavior classification

We further evaluate the ability of the frozen foundation models to perform few-shot behavior classification (Table 3). Similar to classification in the previous section, we use a trainable pooling layer followed by a linear classifier to map model features to class labels, with the difference that the training set only contains a small number of annotated examples per class. These training sets are uniformly randomly sampled for each behavior class from the full set. Results show that VideoPrism outperforms CLIP on all amounts of training data. Notably, VideoPrism achieves results comparable to both the domain-specific baseline and CLIP (Table 2) on Calico (trained on 1, 253 clips) with only 10-shot learning. On both datasets, VideoPrism at 10-shot outperforms CLIP at 100-shot, and further improves as the number of training samples increase. Our training set is randomly sampled to perform this evaluation, and future research integrating foundation models with active learning techniques holds the potential to further improve few-shot performance.

**Table 3.** Few-shot behavior classification results. We use mean average precision over all classes, with the same evaluation setup as Section 2.1. Here, the training set is sampled to 10-shot, 50-shot and 100-shot. Mean and standard deviations of the results are reported over 3 random samples of the training set.

| Dataset | Method | 10-shot | 50-shot | 100-shot |
| --- | --- | --- | --- | --- |
| Calico | CLIP | $39.8 \pm 2.1$ | $42.6 \pm 1.0$ | $47.5 \pm 0.1$ |
| | VideoPrism | $50.5 \pm 3.9$ | $66.3 \pm 0.9$ | $60.2 \pm 0.5$ |
| CalMS21 | CLIP | $27.5 \pm 4.7$ | $42.0 \pm 1.8$ | $49.1 \pm 4.0$ |
| | VideoPrism | $55.5 \pm 3.1$ | $69.8 \pm 2.5$ | $75.0 \pm 2.2$ |

### 2.3 Video-based behavior retrieval

Given a query video from a user, the goal of video-based retrieval is to find similar videos in a large index set of other videos. Retrieval enables researchers to perform targeted search for specific behaviors or events, and rapidly identify relevant footage within large video archives. This capability is particularly crucial in the study of rare behaviors, where manual review to find similar videos over hundreds of hours of data would be prohibitively time-consuming. To assess the potential of foundation models for this task, we investigate their performance in nearest-neighbor retrieval of similar videos using frozen embeddings, without any downstream adaptation.

We present video-based retrieval results on Calico and CalMS21 (Table 4). We visualize retrieval examples (Figure 4, Figure 5) as well as evaluate the models quantitatively using mean Hit@*K*. Hit@*K* is a common metric used to evaluate how accurate the retrievals match the class of the query (over *K* retrievals). Here we use mean Hit@*K*, which is the Hit@*K* averaged over each class of interest to account for class imbalance. We use the test set as the query set, and the train set as the index. We apply global average pooling over spatiotemporal features from both models into a single embedding vector for nearest-neighbor retrieval. On both datasets, we find that VideoPrism features outperform CLIP, and with a high probability of retrieving a video with the same behavior-of-interest with just 5 to 10 retrieval samples. This is especially important for rare behaviors (e.g., “attack” in CalMS21 which only occurs in 2.8% of the frames), suggesting the potential of foundation models for finding videos with similar behaviors in large index sets. However, we observe that in datasets with lighting variations (Figure 4), this retrieval process may prioritize examples with similar lighting conditions alongside similar behaviors. A potential mitigation strategy could involve data augmentation on both the query and index set. This highlights opportunities for further improvements in video retrieval systems using foundation models.

**Fig. 4.**
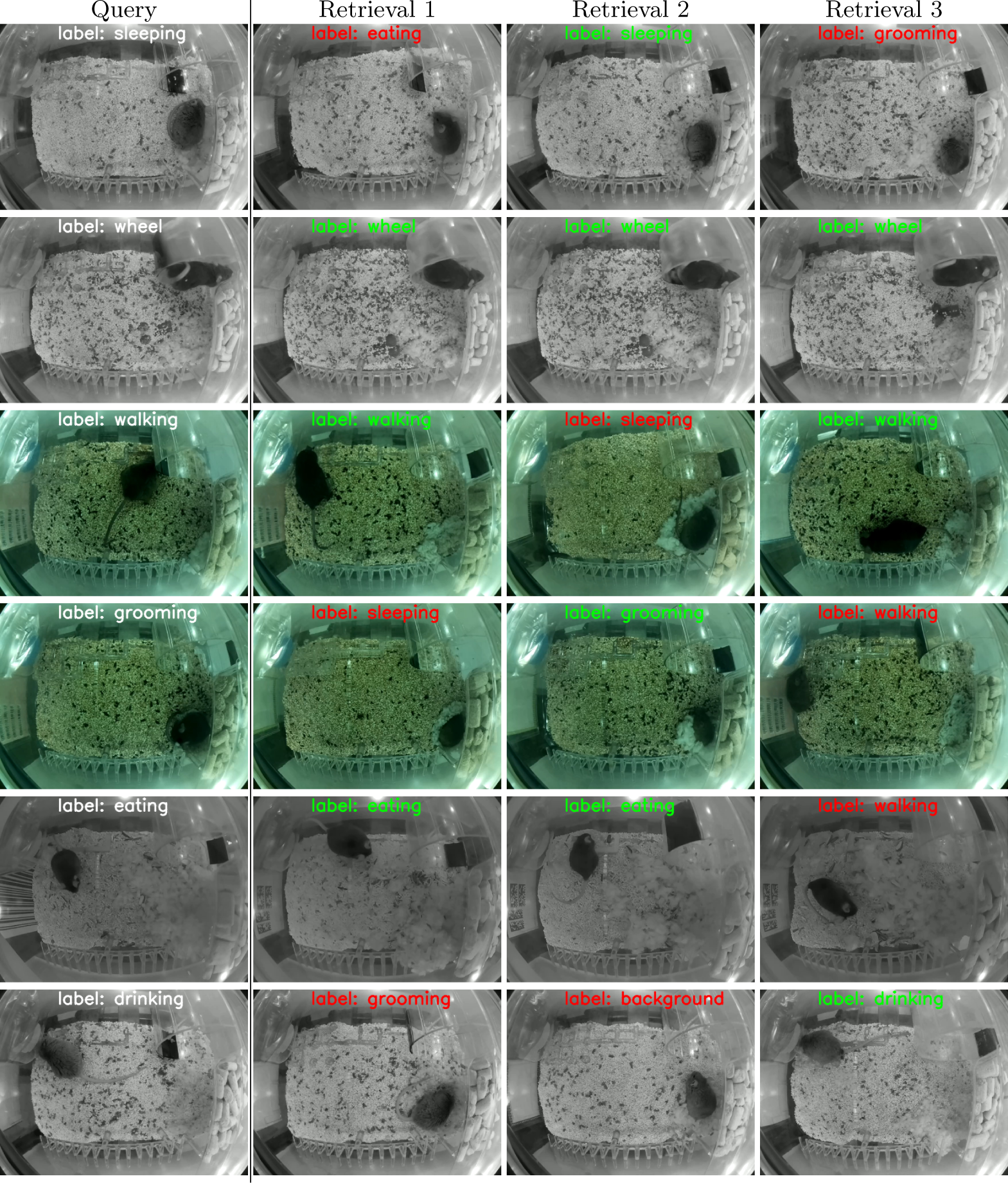
Qualitative video-based retrieval results on Calico. The first column in each row corresponds to an query video (sampled from the test set), and the second to last column corresponds to the three nearest neighbor retrieval results using VideoPrism features on the index set. The expert-annotated behaviors are visualized on each video to show correctness (whether the annotations of the retrieved video matches the query).

**Fig. 5.**
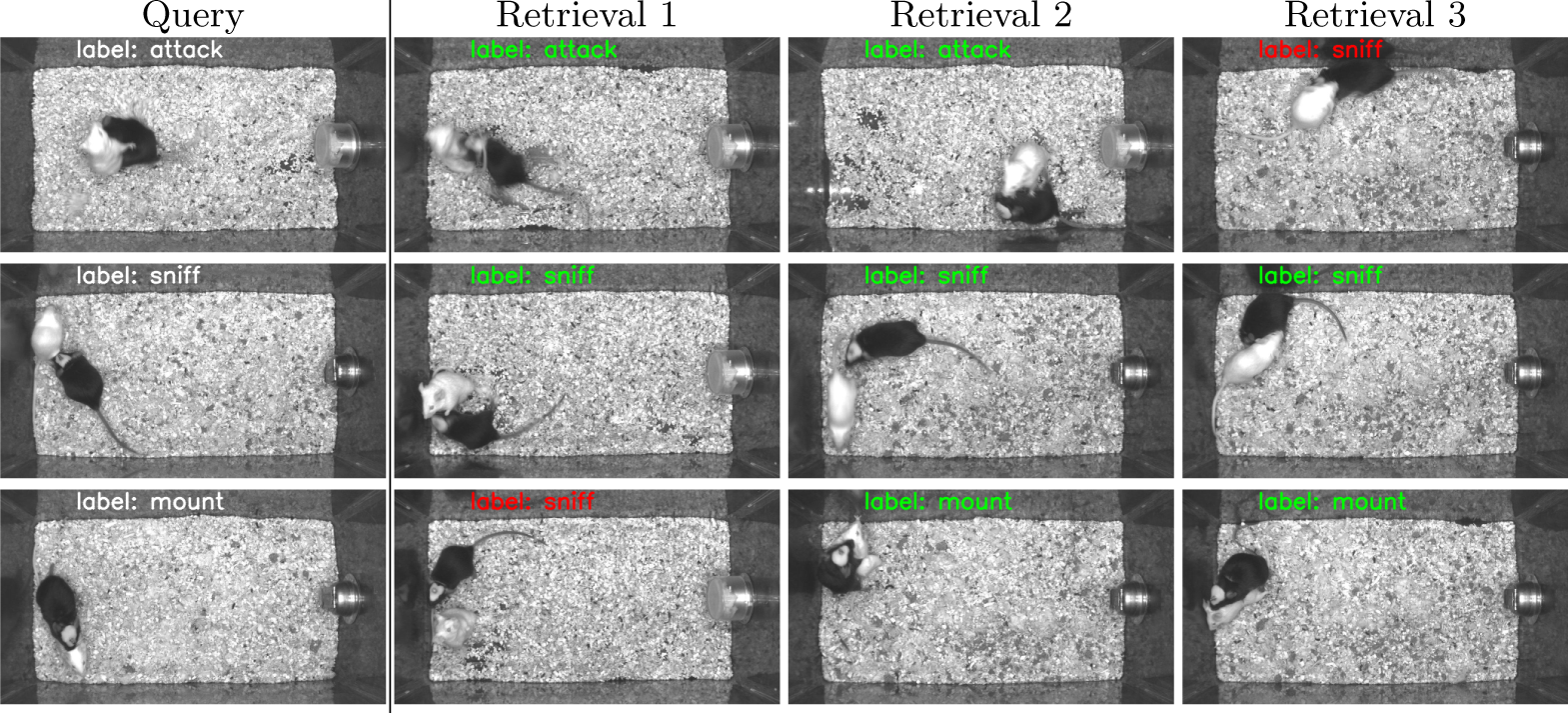
Qualitative video-based retrieval results on CalMS21. The first column in each row corresponds to an query video (sampled from the test set), and the second to last column corresponds to the three nearest neighbor retrieval results using VideoPrism features on the index set. The expert-annotated behaviors are visualized on each video to show correctness (whether the annotations of the retrieved video matches the query).

**Table 4.** Video-based behavior retrieval results. We use mean Hit@*K* averaged over each of the classes (except for the background class “other” in CalMS21, consistent with the behavior classification experiments), computed using the test set as the query set, and the train set as the index set. Random corresponds to randomly retrieving examples from the index set.

| Dataset | Method | mean Hit@1 | mean Hit@5 | mean Hit@10 |
| --- | --- | --- | --- | --- |
| Calico | Random | 16.0 | 47.1 | 65.8 |
|  | CLIP | 28.4 | 58.4 | 75.6 |
|  | VideoPrism | 36.4 | 73.6 | 85.8 |
| CalMS21 | Random | 19.7 | 47.4 | 62.2 |
|  | CLIP | 28.5 | 56.1 | 69.9 |
|  | VideoPrism | 42.0 | 66.2 | 75.1 |

### 2.4 Localization

To study the applicability of foundation models to spatial vision tasks, we assess their performance in localizing animal agents. We present localization results on a video-based mouse dataset Calico-Detection [55] and an image-based mouse dataset MARS [5] ^1^ (Table 5). Calico contains annotated videos (12 frames each) with bounding boxes annotated for each frame, containing a single mouse, while MARS contains one black mouse and one white mouse with two views (top and side). VideoPrism is designed to take a video clip of 16 frames as input, and for the image-based dataset, we repeat the image 16 times to form the input video clip. For evaluation, we follow the standard COCO detection protocol [56] and report the mean average precision over multiple intersection over union thresholds (mAP), in addition to the average precision under 50% and 75% intersection over union (AP@50IoU and AP@75IoU).

**Table 5.** Mouse localization results. We follow the standard COCO detection metrics and report the mean average precision over multiple intersection-over-unions (mAP). We also highlight the APs over 50% and 75% IoU.

| Dataset | Method | mAP | AP@50IoU | AP@75IoU |
| --- | --- | --- | --- | --- |
| Calico-Detection | CLIP | 94.0 | 98.9 | 97.5 |
|  | VideoPrism | 95.2 | 98.9 | 96.0 |
| MARS-Top | CLIP | 58.2 | 97.9 | 64.6 |
|  | VideoPrism | 61.1 | 98.3 | 71.2 |
| MARS-Side | CLIP | 43.9 | 91.1 | 35.3 |
|  | VideoPrism | 46.3 | 92.5 | 40.5 |

Results demonstrate that VideoPrism outperforms CLIP under mAP metric on all datasets (Calico-Detection, MARS-Top, and MARS-Side). Both VideoPrism and CLIP achieve a high performance for AP under 50%IoU, which means that features from both models can roughly locate the mouse; however for AP under 75% IoU, VideoPrism generally performs better than CLIP, which indicates VideoPrism produces more accurate bounding boxes compared to CLIP. For qualitative results (Figure 6), we find that on Calico-Detection, where there is only one mouse, both VideoPrism and CLIP work well. However, on MARS-Top and MARS-Side, which have two mice in each frame, the bounding boxes from VideoPrism is more accurate. Similar results can be observed from Figure 7, in which we compare the precision-recall curves of VideoPrism and CLIP model on these datasets.

**Fig. 6.**
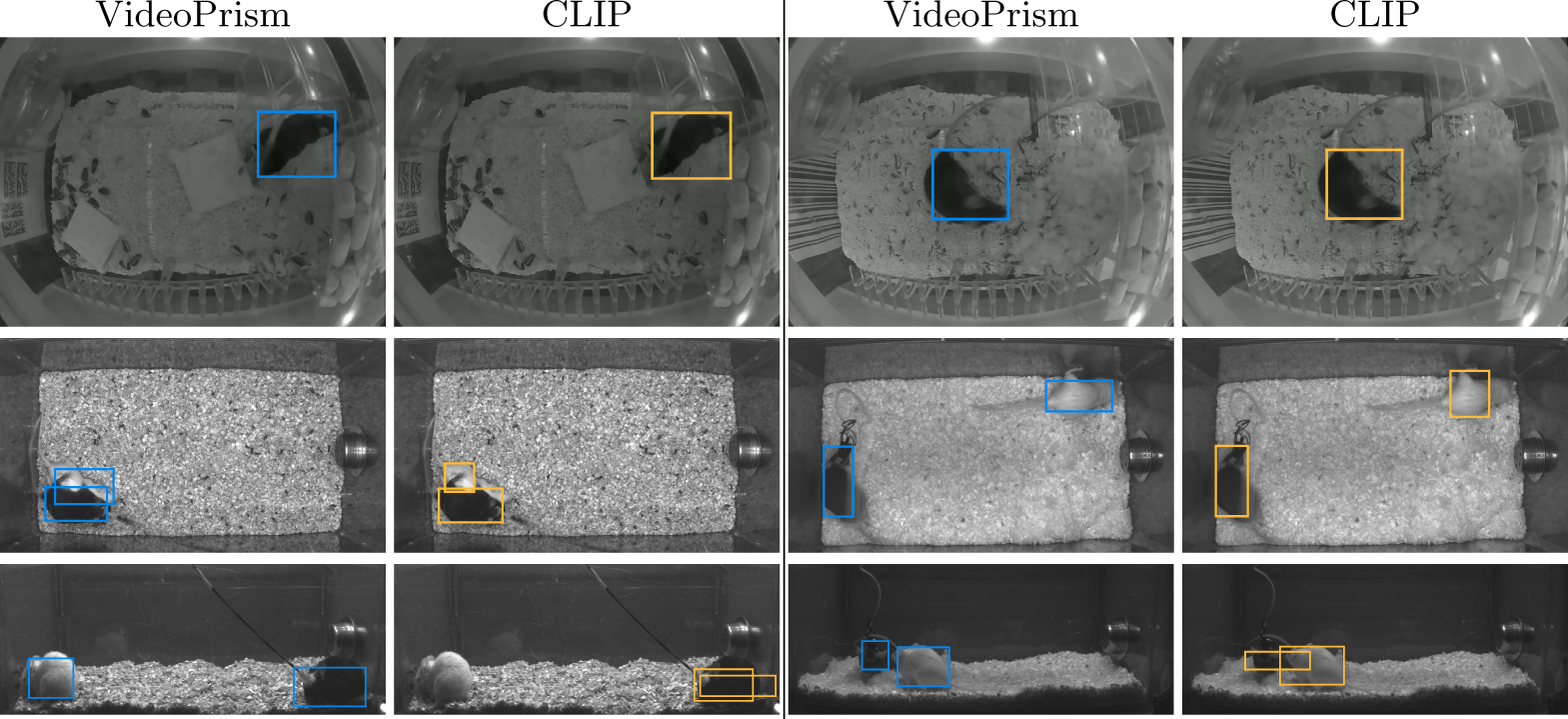
Qualitative comparison of VideoPrism and CLIP on localization. Each row shows localization examples from VideoPrism and CLIP. First row is on the Calico-Detection dataset, second row shows results on MARS-T and third row shows results on MARS-S.

**Fig. 7.**
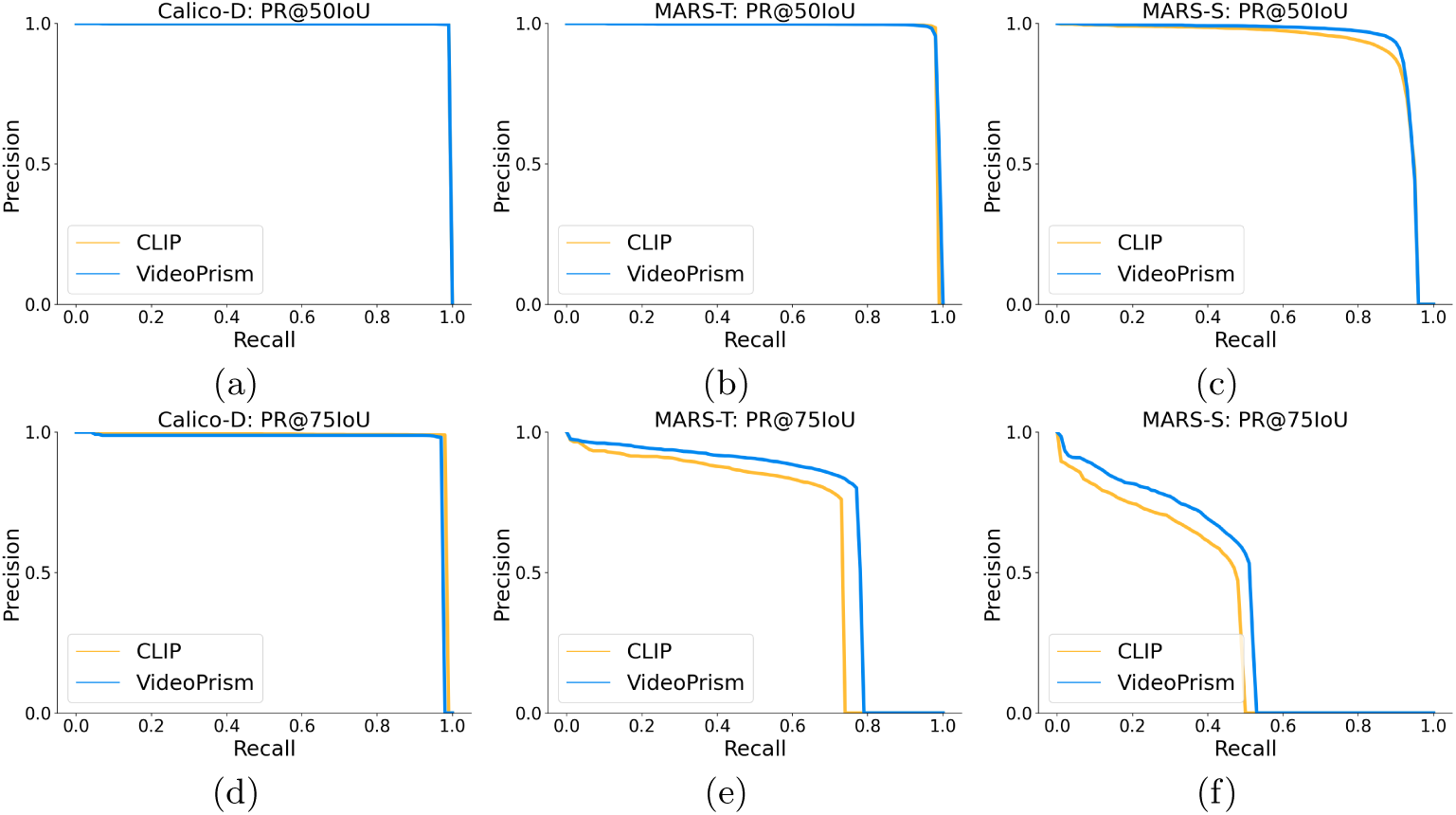
Precision-recall curves for localization. (a) (b) and (c) show the precision recall curve at 50% IoU on Calico, MARS-T and MARS-S dataset. (d) (e) and (f) show the precision recall curve at 75% IoU on Calico, MARS-T and MARS-S dataset.

### 2.5 Broader applications

To evaluate the generalizability of foundation models across diverse species and domains, we evaluate performance on three distinct video-based animal datasets captured across different fields of science: Fly vs. Fly (fly behavior classification), KABR (ecological behavior classification of zebras and giraffes), and SSW60 (fine-grained bird species classification) (Table 6).

**Table 6.** Broader application results. We evaluate model performances using the metrics defined in the original dataset papers. For Fly vs. Fly [19], we use the split in [11] with the mAP metric for behavior classification. For SSW60 [20], we use accuracy as the dataset is balanced. For KABR [21], we use macro-accuracy averaged over classes.

| Method | Fly vs. Fly | SSW60 | KABR |
| --- | --- | --- | --- |
| Specialized models | 88.6 | 71.9 | 61.9 |
| CLIP | 61.7 | 48.3 | 32.2 |
| VideoPrism | 89.1 | 70.1 | 61.6 |

Across all datasets, VideoPrism outperforms CLIP and achieves comparable or better performance to specialized baselines. In the Fly vs. Fly dataset [19], the specialized baseline is a model that leverages preprocessed trajectory information and hand-crafted features [11]. In SSW60 [20], the specialized model is a ResNet-50 pre-trained on iNaturalist and fine-tuned end-to-end on the train split of SSW60. On KABR, the specialized model is a X3D network [57] trained end-to-end on the train split. Each of these specialized models requires either specialized model design, or fine-tuning of the entire pipeline. In comparison, the backbones of both CLIP and VideoPrism are frozen across all the settings. While CLIP does not perform as well as specialized models, VideoPrism is able to achieve comparable performance across species and behaviors, demonstrating its potential to streamline animal behavior research in various ecological and experimental contexts.

## 3 Discussion

The widespread adoption of video technology in animal behavior research has resulted in an explosion of available data, yet the extraction of meaningful insights from these vast video archives remains a significant bottleneck. This usually requires a team of computer vision and machine learning experts to develop a bespoke model for a specific task, under a special setting. Our findings suggest a paradigm shift in this landscape. We demonstrate the generalizability of frozen video foundation models across diverse datasets and tasks and highlight the potential of using a single set of general-purpose video features for animal behavior analysis, without data-specific fine-tuning. This could empower researchers from various disciplines to readily leverage these powerful models, potentially accelerating discoveries in ethology, neuroscience, ecology, and conservation biology.

### 3.1 In-context learning with a general multi-modal LLM

State-of-the-art multi-modal large language models (LLMs) have also been applied to video understanding tasks [3, 58]. Long context windows of such models allow incontext learning, where video frames and associated labels are provided in the prompt. However, the high dimensionality of video data currently requires significant temporal downsampling, limiting their applicability to analyses requiring fine-grained temporal resolution, such as discerning behaviors involving rapid movements or localization.

Despite this limitation, the integration of video foundation models with multi-modal LLMs holds significant promise for unlocking a new paradigm of in-context learning for animal behavior analysis. By leveraging the LLMs’ capacity to process natural language instructions alongside the video foundation model’s ability to extract meaningful representations from video data, we envision a system capable of rapidly adapting to novel behaviors or species without the need for costly and time-consuming retraining or annotation.

Researchers could define new behaviors using simple language descriptions and provide a few illustrative video examples. The multi-modal LLM would then generate task-specific embeddings based on both the language and video input, effectively “tuning” the video foundation model on-the-fly. This approach could empower scientists to explore a wider range of behaviors with minimal effort, significantly accelerating research across fields. Furthermore, integrating data from diverse sources, such as species descriptions, ecological metadata, and scientific literature, could enhance the model’s contextual understanding, leading to more accurate and nuanced interpretations of animal behaviors.

### 3.2 Model cost and latency

The foundation models used in this work currently require considerable memory and computing power to run, and are not yet able to support real-time inference on general hardwares. There are a multitude of ways to alleviate both issues. One way that researchers have studied is to perform model distillation [59]. This approach aims to distill larger models using domain-specific data, and produce a compact model that is economical to run. Additionally, while the foundation models studied in this paper use standard Vision Transformer [60], efficient vision models, such as [61–63], that significantly reduce the computational cost while maintaining competitive performances are an active research topic and provide another promising option for cost reduction.

### 3.3 Role of behavioral video datasets

The success and continued advancement of video foundation models for behavior quantification are closely linked to the availability of diverse and well-annotated video datasets. While we have demonstrated the effectiveness of such models on existing datasets, the limited availability of large-scale, well-annotated, open-sourced video data remains a bottleneck. A concerted effort to curate and release open-sourced datasets, encompassing a wide range of species, behaviors, and environmental contexts, would provide invaluable evaluation data for understanding the capabilities and gaps of video foundation models. Additionally, these data could also be part of training, in order to improve their ability to generalize across scenarios and improve their performance on real-world applications.

Open-sourced video datasets also have the potential to revolutionize the broader field of animal behavior research. By fostering collaboration and knowledge exchange among scientists, these datasets would enable the development of standardized annotation protocols, benchmarking tools, and novel analytical approaches. This collective effort would accelerate methodological advancements, improve reproducibility and comparability across studies, and ultimately deepen our understanding of the complex mechanisms underlying animal behaviors. Moreover, open data would democratize access to cutting-edge research, enabling scientists from diverse backgrounds and institutions to contribute to, and benefit from, this exciting field.

### 3.4 Conclusion

In this work, we have demonstrated the effectiveness of frozen video foundation model on a range of animal behavior datasets and tasks. By leveraging recent machine learning developments, we show that a foundation model’s ability to extract general-purpose features from high-dimensional video data could be applied to a range of tasks without task-specific fine-tuning. These models could be used to compute genera-purpose features in a wide range of settings, opening new avenues for research in ethology, neuroscience, ecology, and conservation biology. While our results demonstrate the strong potential of frozen video foundation models for animal behavior analysis, several promising avenues for future research remain. These include integrating video foundation models with LLMs to enable in-context learning, reducing the computational requirements of video foundation models, and expanding their application to broader ethological questions. We envision a future where video foundation models serve as a robust and adaptable backbone for video-based behavior analysis, enabling researchers to address complex biological questions at unprecedented scales and efficiency.

## 4 Method

We use VideoPrism as our pre-trained video foundation model in our experiments. This model is pre-trained on general Internet videos, without any specific focus to animal datasets. The weights of the model are frozen for all our experiments. In Section 4.1, we summarize the pre-training data used by VideoPrism, Section 4.2 summarizes the VideoPrism architecture, Section 4.3 summarizes the VideoPrism pretraining approach. Finally, Section 4.4 describes our downstream adaptation method for classification, few-shot classification, retrieval, and localization.

### 4.1 Pre-training data

We choose VideoPrism [17] as the video foundation model for our experiments in this paper. At its core, VideoPrism is a general-purpose video encoder. It is pre-trained on 618M web video clips from various internal and external sources. Among them, 36M clips are paired with high-quality manually labelled captions, while captions of the rest video clips are noisy parallel text obtained through automatic speech recognition, metadata, and large multi-modal models [64, 65]. These pre-training data offer an effective source for VideoPrism to learn semantic clues on both object motions and appearances in the video domain, which is fundamental to foundation models [2]. It is noteworthy that we consider the training video corpus general-purpose and has significant difference in distribution from the animal behavior video datasets.

### 4.2 Model architecture

The VideoPrism model adopts a spatial-temporal factorized design based on the Vision Transformer [60] architecture following [66]. At inference time, the model takes 16 video frames of size 288 *×* 288, each of which is evenly partitioned into 16 *×* 16 non-overlapping patches (i.e., visual tokens). These patches are first input to the spatial layers of model in a frame-by-frame fashion. Then, the temporal layers are used for capturing visual semantics across patches in the same spatial locations along the time dimension. We note that the VideoPrism model produces spatiotemporal features in the output token sequence, facilitating the downstream tasks that require fine-grained features (e.g., spatiotemporal action localization). VideoPrism offers two model configurations: VideoPrism-g contains 1B parameters in the spatial encoder, and VideoPrism-B is a base-scale variant with the ViT-Base network [67] (with 80M parameters) as the spatial encoder. For the experiments in this paper, we use the base-scale VideoPrism-B model.

### 4.3 Pre-training approach

The VideoPrism model is pre-trained with a two-stage approach including video-text contrastive learning and masked video modeling. In the first stage, videos and their corresponding captions are encoded individually through a video encoder and a text encoder, producing a global embedding for each modality, respectively. The contrastive loss [4, 29] is optimized between the embeddings of video-text pairs so that they are gradually aligned. This stage yields a video encoder capturing rich visual semantics from language supervision, which supplies semantic video embeddings for the second-stage training. In the second-stage, the video encoder is continually trained with only videos based on an improved paradigm of masked video modeling [42, 68, 69]. To be specific, the VideoPrism model is trained to predict the video-level global embedding and also token-wise embeddings from the first-stage model to effectively leverage the knowledge acquired in that stage. Meanwhile, the predicted tokens are randomly shuffled to prevent the model from learning shortcuts. The training of this stage focuses on learning both appearance and motion information from only videos.

### 4.4 Adaptation to downstream tasks

We adapt VideoPrism to four animal behavioral analysis tasks including behavior video classification, few-shot behavior video classification, video-based behavior retrieval, and localization. The pre-trained VideoPrism is used as an encoder to extract features from the input video clips. For all these tasks, the VideoPrism is fixed and only the task-specific decoder is trained to adapt to the task when necessary. We also adapt the frozen image foundation model CLIP in a comparable way, described in more detail below.

#### 4.4.1 Behavior classification

In behavior classification, a model is designed to take a video clip containing animal behavior and classify it into several expert-defined categories. For this task, following [34], a multi-head attention pooling (MAP) layer is employed as the task-specific decoder which plays the role of a classification head. It contains a Transformer layer which takes a trainable query token and the spatiotemporal tokens from VideoPrism as input. The query token aggregates information from the visual tokens through cross attention and a softmax layer takes the aggregated feature as input and predicts the class probability.

To handle videos with an image foundation model (e.g., CLIP), a conventional approach [32] is to apply the image encoder on video frames individually without early fusion of temporal information and append an MAP layer to extract the global representation from concatenated per-frame features. However, this approach is problematic when input videos are motion-heavy, because there are no temporal signals incorporated during the downstream adaptation. To address this issue, we introduce an improved pooling method with a factorized design in space and time. Specifically, we first apply a spatial pooling layer to video frames separately to produce a per-frame global representation for each of them. After adding temporal positional embeddings [22] to all per-frame global embeddings to encode temporal information, we apply a temporal pooling layer to extract the global representation for the entire video. By doing so, we observe a notable improvement over the conventional pooling approach, and thus we use this method in our experiments. We call this method the factorized multi-head attention pooling (FMAP).

**Fig. 8.**
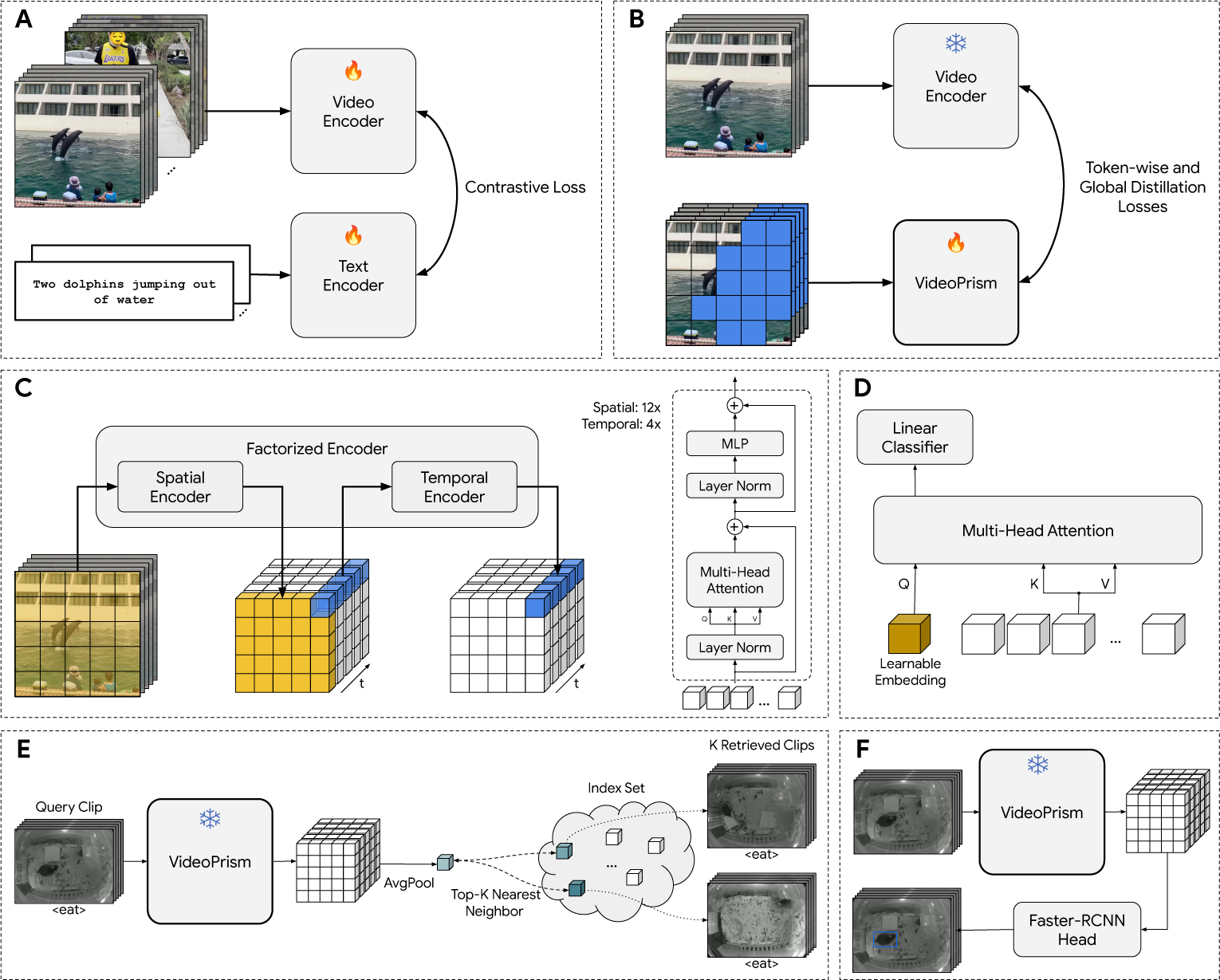
Methods overview. VideoPrism [17] is pre-trained with a two-stage approach. **(A)** In the first stage, a video encoder and a paired text encoder are via contrastive loss over a large amount of video-text pairs. **(B)** In the second stage, VideoPrism is trained via a masked modeling approach with token-wise and global self-distillation losses. **(C)** The VideoPrism model is based on a spatial-temporal factorized encoder architecture [66], which encodes spatial information frame-by-frame, followed by temporal aggregation. The encoder layers use standard self-attention Transformer architecture [70]. **(D)** For downstream classification tasks, we further train multi-head attention pooling layer over frozen spatiotemporal features from a video clip, followed by a linear classifier. **(E)** For video retrieval tasks, we do global average pooling over frozen spatiotemporal features and use the pooled embedding for nearest neighbor retrieval for top-K similar clips. **(F)** For localization tasks, we apply a Faster-RCNN head [71] frame-by-frame on top of frozen spatial-temporally aggregated features for bounding box prediction.

#### 4.4.2 Few-shot behavior classification

Few-shot classification is similar to behavior classification, but only a few labeled examples for each category is used to train the task specific decoder. We use the same MAP layer as the classification method for VideoPrism, and the same FMAP layer as the classification method for CLIP described in Section 4.4.1. Similar to before, a softmax operator is used to output the class probability.

#### 4.4.3 Video-based behavior retrieval

The goal of video-based retrieval is to search through a large database of videos for ones that are similar to the query video. For animal behavior analysis, we would like to search for videos that show similar behaviors as the query. In this task, we use pretrained VideoPrism, without any task-specific decoder, to extract embeddings from all query videos and videos in the index dataset. We first apply VideoPrism to the input video clip to obtain the spatiotemporal tokens for the query, then a non-trainable global average pooling is applied to these tokens to get the final representation embedding for the input video. The most similar videos from the index dataset are selected based on the *L*_2_ distance of their embedding with that of the query video. There is no training for obtaining the retrieval embeddings, and the same setup of average pooling over the tokens is used for CLIP.

#### 4.4.4 Localization

A localization model usually indicates the location of the target objects by predicting the boxes tightly bounding them. To localize animals with VideoPrism, we concatenate a Faster-RCNN [71] detection head as the task-specific decoder to the VideoPrism backbone. Following [72], we take the the features output from VideoPrism and apply convolutions with stride *{*4, 2, 1, 1*/*2, 1*/*4*}* to form the simple feature pyramid. This feature pyramid will then be input to the Faster-RCNN head for predicting bounding boxes. More specifically, to apply detection on the *i*-th frame *f_i_* in a video, we extract 16 frames around the target frame *f_i_*to form the input video clip *{f_i−_*_7_*,…f_i_, …, f_i_*_+8_*}*. After extracting features from VideoPrism, only the features that correspond to the target frame *f_i_* are fed into the Faster-RCNN head to predict bounding boxes in the target frame. To apply detection on a single image, we directly repeat the image 16 times to form a static video clip and only use the feature of the middle frame from VideoPrism for detection. We use CLIP in the same way for the localization tasks.

### 4.5 Experimental setup

We use the following open-sourced, annotated datasets:

- **CalMS21** [18]: CalMS21 Task 1 contains top-view videos capturing mice social behavior in a standard resident-intruder assay at 30Hz. The videos consist of freely interacting mice, annotated with frame-level labels for social behaviors (e.g., “attack”, “mount”, “sniff”). The challenge lies in accurately recognizing and classifying these behaviors from the raw video data.
- **Calico** [55]: Calico consists of annotated video clips of mice in their home cage at 24Hz. Each video clip has one mouse, and is annotated with clip-level labels for common behaviors (e.g., “walking”, “grooming”, “drinking”).
- **Calico-Detection** [55]: Calico-Detection is a part of the Calico dataset that has additional bounding box annotations to study localization methods. There is a bounding box annotated per frame for the mouse.
- **CRIM13** [9]: CRIM13 consists of videos collected in a standard resident-intruder assay at 25Hz. Different from CalMS21, this set includes conditions such as human interventions (e.g., hands reaching into the cage), as well as two camera views (top-view and side-view). The videos are annotated at the frame-level with behaviors-of-interest (e.g., “approach”, “chase”, “drink”). We present results of our models on both CRIM13-Top and CRIM13-Side. In the original paper, CRIM13 contains matching top and side videos, but in the dataset available to download, not all recorded videos match between the top and side views. We report our performance separately on each view.
- **MARS** [5]: We use the MARS dataset for its annotated detection bounding boxes – the original dataset is captured as video, but only non-continuous images are annotated with bounding boxes. There is one box annotated for each of two mice. We evaluate on both the top view and side view of MARS.
- **Fly vs. Fly** [19]: Fly vs. Fly consists of per-frame expert-annotated behaviors of a pair of flies. We use the videos captured at 30Hz that study the effect of genetic manipulation on behaviors including aggression and courtship. The baseline for this model uses trajectory data, and as not all frames of the video contains valid tracks, we use the same data split as [11] for this dataset to match the video and trajectory dataset for evaluation. The dataset contains the following behaviors: “lunge”, “wing” “threat”, “tussle”, “wing” “extension”, “circle”, and “copulation”.
- **KABR** [21]: KABR captures drone videos of Kenyan wildlife, including reticulated giraffes, plains zebras, and Grevy’s zebras. The dataset is captured as sequences of videos containing different individuals of interest, and annotated by ecologists for behaviors such as “graze”, “trot”, and “head up”.
- **SSW60** [20]: SSW60 consists of videos of birds collected from the Sapsucker Woods. The dataset is balanced in terms of classes, with 60 species of birds for fine-grained species classification. As the birds are moving, and not all frames of the video contains the birds, video information is helpful for classification. The dataset additionally includes audio, which we do not use in this evaluation.

To facilitate comparison of the evaluation results, we use the same train and test split of the data as domain-specific baselines, when reproducible splits exist and are available. Notably, the foundation models are not trained on the train split, and require no or minimal adaptation. In tasks where adaptation is required, the train split is used to update downstream task decoders, described in Section 4.4.

## Acknowledgements

We would like to thank Nitesh B. Gundavarapu, Shen Yan, Luke Friedman, Rui Qian, Tobias Weyand, Yue Zhao, Rachel Hornung, Ming-Hsuan Yang, Huisheng Wang, Mikhail Sirotenko, and Boqing Gong for their contribution to the VideoPrism project at Google. We thank Alex Siegman, Amanda Sadler, and Rachel Stigler for their support for this project. We also would like to thank Rahul Sukthankar, Tomas Izo, and Caroline Pantofaru for their leadership support. Additionally, we want to thank Ann Kennedy at Northwestern for helpful discussions on animal behavior. All these contribution and support made this project possible.

## Supplementary: Implementation Details

### Evaluation Setup

We extract frozen features from foundation models, and use this set of features across all datasets. We summarize the domain specific models used in each dataset below. Calico [55] is used for mouse behavior classification from the top view, and the domain expert model [55] is a ResNet encoder with features averaged temporally, trained with a 2-layer multi-layer perceptron on Calico. CalMS21 [18] is used for mouse social behavior video classification from top view, and the domain expert model [18] is trained on a combination of unlabelled and labelled data with hand-crafted trajectory features using task programming [11]. CRIM13 [9] is used for mouse behavior video classification with top and side views, and in the original dataset paper, the domain-specific model combines side and top views. However, in the current open-sourced version, the sequences are mis-matched between the views, and we evaluate on each separately. Fly vs. Fly [19] is used for fly behavior video classification, and the data split and domain expert model is from task programming [11], trained using a combination of hand-crafted features and the annotated training set. KABR [21] is used for for behavior video classification with Kenyan animals, and the domain-specific model is an X3D [57] video architecture trained on KABR. SSW60 [20] is used for fine-grained bird species video classification, and the domain-specific model is a ResNet50 pre-trained on iNaturalist [73] (a dataset for fine-grained animal species classification) and fine-tuned on SSW60.

For localization, Calico-Detection [55] did not provide a domain-specific localization baseline. MARS [5] originally uses a multi-scale convolutional MultiBox [74] approach for detection, but only the train split of the original paper is open-sourced (15k frames). We create our own train/test split from the open-sourced dataset.

For all other datasets, we use the train and test splits defined by existing work when videos are available. Notably, existing work split train and test sets at the video-level (not the frame-level). We use the same metrics as domain-specific models when available, which is mAP for all works, except KABR (macro-accuracy averaged across classes) and SSW60 (accuracy). We also note that following previous work [11, 18], in datasets where there are frame-level annotations including background classes (CalMS21, CRIM13, Fly vs. Fly), the metric is only averaged across behaviors-of-interest (not including background classes).

We extract all frames from the video at the original FPS of each dataset. We use 16 consecutive frames as input for all datasets, and 16 frames with a stride of 5 for KABR (following the baseline).

### Downstream Adaptation

We include more details about each downstream application.

#### Behavior classification

We study behavior classification on four datasets: Calico [55], CalMS21 [18] for mouse video classification from top view, CRIM13 [9] for mouse video classification with top and side views. We follow existing work to split the four datasets into train and test subset. We use mAP as the metric to evaluate the classification performance. For datasets such as CalMS21 [18] which includes background classes, the metric is only averaged across behaviors-of-interest without background class.

When training behavior classification, we follow the VideoGLUE [34] frozen-backbone setup which is also adapted in VideoPrism [17] for downstream tasks. To train the classification head, we employ the cross entropy loss defined as follows:

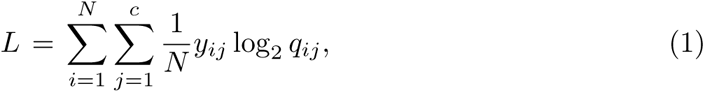

where *N* is the number of training samples and *c* is the number of behavior categories. *y_ij_ ∈ {*0, 1*}* represents the ground truth class label for each sample and *q_ij_* represents the predicted probability of the *i*-th sample being assigned to *j*-the class through our classification model. We use AdamW [75] as the optimizer with a cosine learning rate decay. For all the experiments, we set the learning rate to be 5*e^−^*^5^ and use dropout rate 0.5.

#### Few-shot behavior classification

Few-shot behavior classification has the same setup as behavior classification, except the training set is sampled. Following the common few-shot setup, we sample 10, 50, and 100-shot per class in each dataset. We perform three random sets of samples, and apply them on each of the models to compute mean and variance. The implementation details are the same as for behavior classification.

#### Behavior retrieval

Retrieval does not require any training, and we simply take the average of spatiotemporal tokens across all frames. We then perform nearest neighbor retrieval based on *l*2 distance between features computed for each clip. We use 16 frames for each of the query and index clips. On Calico, we use the full train and test split as the index and query. On CalMS21, we remove the background class “other” from the query set, since we want to focus on retrieving the behaviors-of-interest (the background class is not removed from the index set). We then perform three runs at 10% subsampling to obtain the average performance using mean Hit@*K*.

#### Localization

For Calico dataset, since there is one mouse in each video, the localization is formulated as single class object detection problem, *i.e.*, each predicted boxes is assigned with either “Mouse” or “Background”. For MARS-Top and MARS-Side dataset, we treat “Black mouse” and “White mouse” as different classes, as a results, localization on these two datasets are formulated as a multiple class object detection problem, each predicted box will be assigned to “Black mouse”, “White mouse” or “Background”. Following general object detection tasks, our loss is defined to have a classification loss and regression loss as follows:

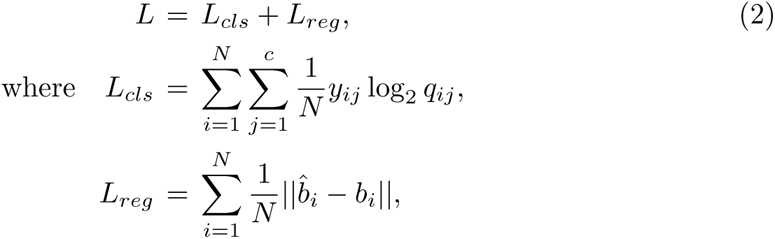

where *L_cls_* is a cross entropy loss used to supervise the predicted class. *N* is the number of samples and *c* is the number of classes (2 for Calico dataset and 3 for MARS-Top and MARS-Side dataset). *y_ij_ ∈ {*0, 1*}* represents the ground truth class label for each sample and *q_ij_* represents the probability of the *i*-th sample being assigned to *j*-the class. *L_reg_* is a *L*1 regression loss used to supervise the predicted bounding box coordinate. ^^^*b_i_* and *b_i_* are predicted and ground truth bounding boxes. We use AdamW [75] optimizer to train our localization model for 3, 000 epochs on Calico dataset and 300 epochs for MARS-Top and MARS-Side dataset. During training, we use the first 10% epochs as the warm up steps in which the learning rate linearly increasing from 0 to 1*e^−^*^4^. After that, the learning rate is decayed to 0 following the cosine annealing schedule. We set the weight decay as 1*e^−^*^5^ through out the training process. Following their pre-training strategy, the input resolution to our localization model with VideoPrism and CLIP backbone are 228 *×* 228 and 224 *×* 224 respectively. For Calico dataset, during training, no data augmentation is used, we direclty pad and resize the video to the target resolution. For MARS-Top and MARS-Side dataset, we apply large-scale jittering [76] and random shift the image horizontaly and vertically as the data augmentation. We use batch size 64 during training.

#### Broader applications

The datasets for broader applications are Fly vs. Fly, SSW60, and KABR. All models are trained with the same implementation details as described in behavior classification. We note that the exception is KABR, which is trained with the EQL loss [77], with 5*e^−^*^6^ learning rate, following baseline to account for class imbalance.

## Footnotes

1 The original CRIM13 study proposed a specialized model using multi-view videos [9]. However, the current public release have mis-matched sequences in each view. Therefore, we evaluate separately in each view instead of combining the views.

1 The original MARS dataset is video-based, however, the authors of [5] only open sourced selected frames from the original video dataset.

## References

[1] Anderson, D. J. & Perona, P. Toward a science of computational ethology. Neuron 84, 18–31 (2014).

[2] Bommasani, R., et al. On the opportunities and risks of foundation models. arXiv preprint arXiv:2108.07258 (2021).

[3] Achiam, J. et al. GPT-4 technical report. arXiv preprint arXiv:2303.08774 (2023).

[4] Radford, A., et al. Learning transferable visual models from natural language supervision. ICML (2021).

[5] Segalin, C. et al. The mouse action recognition system (MARS) software pipeline for automated analysis of social behaviors in mice. Elife 10, e63720 (2021).

[6] Luxem, K. et al. Open-source tools for behavioral video analysis: setup, methods, and best practices. elife 12, e79305 (2023).

[7] Nilsson, S. R., et al. Simple behavioral analysis (SimBA): an open source toolkit for computer classification of complex social behaviors in experimental animals. BioRxiv (2020).

[8] Hong, W. et al. Automated measurement of mouse social behaviors using depth sensing, video tracking, and machine learning. Proceedings of the National Academy of Sciences 112, E5351–E5360 (2015).

[9] Burgos-Artizzu, X. P., Dollár, P., Lin, D., Anderson, D. J. & Perona, P. Social behavior recognition in continuous video. IEEE Conf. Comput. Vis. Pattern Recog. (2012).

[10] Graving, J. M. et al. Deepposekit, a software toolkit for fast and robust animal pose estimation using deep learning. Elife 8, e47994 (2019).

[11] Sun, J. J., et al. Task programming: Learning data efficient behavior representations. IEEE Conf. Comput. Vis. Pattern Recog. (2021).

[12] He, K., Zhang, X., Ren, S. & Sun, J. Deep residual learning for image recognition. Proceedings of the IEEE conference on computer vision and pattern recognition 770–778 (2016).

[13] Deng, J. et al. ImageNet: A large-scale hierarchical image database. 2009 IEEE conference on computer vision and pattern recognition 248–255 (2009).

[14] Bohnslav, J. P. et al. DeepEthogram, a machine learning pipeline for supervised behavior classification from raw pixels. Elife 10, e63377 (2021).

[15] Mathis, A. et al. DeepLabCut: markerless pose estimation of user-defined body parts with deep learning. Nature neuroscience 21, 1281–1289 (2018).

[16] Pereira, T. D., et al. Sleap: A deep learning system for multi-animal pose tracking. Nature methods 19, 486–495 (2022).

[17] Zhao, L. et al. VideoPrism: A foundational visual encoder for video understanding. ICML (2024).

[18] Sun, J. J., et al. The multi-agent behavior dataset: Mouse dyadic social interactions. NeurIPS datasets and benchmarks (2021).

[19] Eyjolfsdottir, E., et al. Detecting social actions of fruit flies. Eur. Conf. Comput. Vis. (2014).

[20] Van Horn, G., et al. Exploring fine-grained audiovisual categorization with the SSW60 dataset. Eur. Conf. Comput. Vis. (2022).

[21] Kholiavchenko, M., et al. KABR: In-situ dataset for kenyan animal behavior recognition from drone videos. WACV (2024).

[22] Devlin, J., Chang, M.-W., Lee, K. & Toutanova, K. BERT: Pre-training of deep bidirectional transformers for language understanding. NAACL (2019).

[23] Brown, T. et al. Language models are few-shot learners. Adv. Neural Inform. Process. Syst. (2020).

[24] Yuan, L., et al. Florence: A new foundation model for computer vision. arXiv preprint arXiv:2111.11432 (2021).

[25] Wang, J. et al. OmniVL: One foundation model for image-language and video-language tasks. Adv. Neural Inform. Process. Syst. (2022).

[26] Xu, H. et al. mPLUG-2: A modularized multi-modal foundation model across text, image and video. ICML (2023).

[27] Girdhar, R., et al. ImageBind: One embedding space to bind them all. IEEE Conf. Comput. Vis. Pattern Recog. (2023).

[28] Zhu, B., et al. LanguageBind: Extending video-language pretraining to N-modality by language-based semantic alignment. Int. Conf. Learn. Represent. (2024).

[29] Jia, C., et al. Scaling up visual and vision-language representation learning with noisy text supervision. ICML (2021).

[30] Chen, X., et al. PaLI: A jointly-scaled multilingual language-image model. Int. Conf. Learn. Represent. (2023).

[31] Alayrac, J.-B. et al. Flamingo: A visual language model for few-shot learning. Adv. Neural Inform. Process. Syst. (2022).

[32] Yu, J. et al. CoCa: Contrastive captioners are image-text foundation models. TMLR (2022).

[33] Singh, A., et al. FLAVA: A foundational language and vision alignment model. IEEE Conf. Comput. Vis. Pattern Recog. (2022).

[34] Yuan, L. et al. VideoGLUE: Video general understanding evaluation of foundation models. arXiv preprint arXiv:2307.03166 (2023).

[35] Gu, X., Lin, T.-Y., Kuo, W. & Cui, Y. Open-vocabulary object detection via vision and language knowledge distillation. ICLR (2022).

[36] Ghiasi, G., Gu, X., Cui, Y. & Lin, T.-Y. Scaling open-vocabulary image segmentation with image-level labels. Eur. Conf. Comput. Vis. 540–557 (2022).

[37] Luo, H., et al. CLIP4Clip: An empirical study of CLIP for end to end video clip retrieval. arXiv preprint arXiv:2104.08860 373 (2021).

[38] Qian, R., et al. Spatiotemporal contrastive video representation learning. IEEE Conf. Comput. Vis. Pattern Recog. (2021).

[39] Feichtenhofer, C., Fan, H., Xiong, B., Girshick, R. & He, K. A large-scale study on unsupervised spatiotemporal representation learning. IEEE Conf. Comput. Vis. Pattern Recog. (2021).

[40] Recasens, A., et al. Broaden your views for self-supervised video learning. Int. Conf. Comput. Vis. (2021).

[41] Singh, A., et al. Semi-supervised action recognition with temporal contrastive learning. IEEE Conf. Comput. Vis. Pattern Recog. (2021).

[42] Wei, C. et al. Masked feature prediction for self-supervised visual pre-training. IEEE Conf. Comput. Vis. Pattern Recog. (2022).

[43] Yuan, L., et al. Contextualized spatio-temporal contrastive learning with self-supervision. IEEE Conf. Comput. Vis. Pattern Recog. (2022).

[44] Qian, R., et al. On temporal granularity in self-supervised video representation learning. Brit. Mach. Vis. Conf. (2022).

[45] Tong, Z., Song, Y., Wang, J. & Wang, L. VideoMAE: Masked autoencoders are data-efficient learners for self-supervised video pre-training. Adv. Neural Inform. Process. Syst. (2022).

[46] Wang, L., et al. VideoMAE v2: Scaling video masked autoencoders with dual masking. IEEE Conf. Comput. Vis. Pattern Recog. (2023).

[47] Zellers, R., et al. MERLOT: Multimodal neural script knowledge models. Adv. Neural Inform. Process. Syst. (2021).

[48] Fu, T.-J., et al. VIOLET: End-to-end video-language transformers with masked visual-token modeling. arXiv preprint arXiv:2111.12681 (2021).

[49] Li, L., et al. LAVENDER: Unifying video-language understanding as masked language modeling. IEEE Conf. Comput. Vis. Pattern Recog. (2023).

[50] Wang, J., et al. All in one: Exploring unified video-language pre-training. IEEE Conf. Comput. Vis. Pattern Recog. (2023).

[51] Cheng, F., et al. VindLU: A recipe for effective video-and-language pretraining. IEEE Conf. Comput. Vis. Pattern Recog. (2023).

[52] Piergiovanni, A., et al. Mirasol3B: A multimodal autoregressive model for time-aligned and contextual modalities. arXiv preprint arXiv:2311.05698 (2023).

[53] Xiong, Y., et al. Spatiotemporally discriminative video-language pre-training with text grounding. arXiv preprint arXiv:2303.16341 (2023).

[54] Wang, Z. et al. Paxion: Patching action knowledge in video-language foundation models. Adv. Neural Inform. Process. Syst. (2023).

[55] Hu, B., et al. 3D mouse pose from single-view video and a new dataset. Scientific Reports 13, 13554 (2023). URL https://www.nature.com/articles/s41598-023-40738-w. Number: 1 Publisher: Nature Publishing Group.

[56] Lin, T.-Y. et al. Microsoft COCO: Common objects in context. Computer Vision– ECCV 2014: 13th European Conference, Zurich, Switzerland, September 6-12, 2014, Proceedings, Part V 13 740–755 (2014).

[57] Feichtenhofer, C. X3D: Expanding architectures for efficient video recognition. Proceedings of the IEEE/CVF conference on computer vision and pattern recognition 203–213 (2020).

[58] Team, G. et al. Gemini: A family of highly capable multimodal models. arXiv preprint arXiv:2312.11805 (2023).

[59] Yao, Y., Huang, S., Wang, W., Dong, L. & Wei, F. Adapt-and-distill: Developing small, fast and effective pretrained language models for domains. URL http://arxiv.org/abs/2106.13474.2106.13474[cs].

[60] Dosovitskiy, A., et al. An image is worth 16x16 words: Transformers for image recognition at scale. Int. Conf. Learn. Represent. (2021).

[61] Piergiovanni, A., Kuo, W. & Angelova, A. Rethinking video ViTs: Sparse video tubes for joint image and video learning. Proceedings of the IEEE/CVF Conference on Computer Vision and Pattern Recognition 2214–2224 (2023).

[62] Yang, T. et al. AIM: Adapting image models for efficient video action recognition. arXiv preprint arXiv:2302.03024 (2023).

[63] Qin, D., et al. MobileNetV4–universal models for the mobile ecosystem. arXiv preprint arXiv:2404.10518 (2024).

[64] Wang, Y., et al. InternVid: A large-scale video-text dataset for multimodal understanding and generation. arXiv preprint arXiv:2307.06942 (2023).

[65] Zhao, Y., et al. Distilling vision-language models on millions of videos. arXiv preprint arXiv:2401.06129 (2024).

[66] Arnab, A., et al. ViViT: A video vision transformer. Int. Conf. Comput. Vis. (2021).

[67] Zhai, X., Kolesnikov, A., Houlsby, N. & Beyer, L. Scaling vision transformers. IEEE Conf. Comput. Vis. Pattern Recog. (2022).

[68] He, K., et al. Masked autoencoders are scalable vision learners. IEEE Conf. Comput. Vis. Pattern Recog. (2022).

[69] Feichtenhofer, C., Fan, H., Li, Y. & He, K. Masked autoencoders as spatiotemporal learners. Adv. Neural Inform. Process. Syst. (2022).

[70] Vaswani, A., et al. Attention is all you need. NIPS (2017).

[71] Ren, S., He, K., Girshick, R. & Sun, J. Faster R-CNN: Towards real-time object detection with region proposal networks. Adv. Neural Inform. Process. Syst. (2015).

[72] Li, Y., Mao, H., Girshick, R. & He, K. Exploring plain vision transformer backbones for object detection. ECCV (2022).

[73] Van Horn, G. et al. The iNaturalist species classification and detection dataset. Proceedings of the IEEE conference on computer vision and pattern recognition 8769–8778 (2018).

[74] Szegedy, C., Reed, S., Erhan, D., Anguelov, D. & Ioffe, S. Scalable, high-quality object detection. arXiv preprint arXiv:1412.1441 (2014).

[75] Loshchilov, I. & Hutter, F. Decoupled weight decay regularization. Int. Conf. Learn. Represent. (2019).

[76] Ghiasi, G. et al. Simple copy-paste is a strong data augmentation method for instance segmentation. CVPR (2021).

[77] Tan, J., et al. Equalization loss for long-tailed object recognition. IEEE Conf. Comput. Vis. Pattern Recog. (2020).

